# Follistatin-like 1 Promotes Neutrophil-mediated Host Defense against *Klebsiella pneumoniae*

**DOI:** 10.64898/2026.10.27.762279

**Authors:** Norie Sugitani, Matthew Henkel, Alexander Applegate, Jessica Partyka, Tianyi Wu, Nathan W. Froelich, Angelika Dewicki, Brian T. Campfield

## Abstract

*Klebsiella pneumoniae* is a leading cause of pneumonia and highly burdensome etiology of antibiotic-resistant infections constituting an urgent public health threat. Follistatin-like 1 (FSTL-1) is implicated in many processes including lung development and homeostasis, tissue remodeling, and inflammation though its role in antibacterial immunity remains poorly defined. We have observed that FSTL-1 is elevated in the lungs of both humans and mice during bacterial pneumonia. Using an inducible *Fstl1* knockout mouse model, we demonstrate that FSTL-1 is required for pulmonary bacterial clearance and neutrophil recruitment during *K. pneumoniae* infection. Recombinant FSTL-1 rescues *Fstl1*-deficient susceptibility and enhances clearance of *K. pneumoniae* in wild-type mice. Mechanistically, FSTL-1-mediated protection was lost upon neutrophil depletion and in *Cd14*-deficient or *Nr4a1* deficient mice – components of a lung-specific neutrophil transcriptional program. These findings identify FSTL-1 as a lung-derived factor that licenses neutrophil-mediated antibacterial defense via a novel FSTL-1/CD14/NR4A1-dependent signaling pathway critical for protection against bacterial pneumonia.

## Introduction

Pneumonia is the leading infectious cause of death among children worldwide, accounting for 15% of overall childhood mortality and surpassed only by prematurity^1,2^. In the US, it is the most common infectious cause of hospitalization and death, with annual costs exceeding $17 billion^3-6^. Although antibiotics remain the standard treatment for bacterial pneumonia^7^, antibiotic resistance (AR),^8-11^ is increasingly common, and AR pathogens are associated with higher costs, morbidity and mortality^12-14^. *Klebsiella pneumoniae* is a leading cause of hospital-acquired pneumonia, constituting an urgent threat to public health owing to its high virulence and rising rates of antibiotic resistance^14^. Improving treatment strategies for AR pneumonia is therefore an urgent need. Because the pathophysiology of bacterial pneumonia is governed by complex host-pathogen interactions^7^, defining the critical mediators of host immunity offers the potential for developing immune-targeted therapies.

Neutrophils are essential effectors of pulmonary host defense^15^. Recent studies have established a revised paradigm where neutrophils are a dynamic population, rather than a classical understanding as a short-lived monotonous population of circulating effector cells; unlike in the bone marrow and blood circulation, neutrophils acquire distinct tissue-specific phenotypes upon entering peripheral organs^16,17^. In the lungs, neutrophils rapidly adopt a specialized transcriptional program – marked by expression of CD14 and NR4A1^16^, that may support effective antibacterial function. The tissue-specific signals that define the lung niche, along with how neutrophils recognize them and adopt a lung-specific neutrophil phenotype remains poorly understood.

Follistatin-like 1 (FSTL-1) is a glycoprotein secreted by mesenchymal and stromal cells that regulates tissue remodeling and development through modulation of BMP4 and TGF-β signaling^18-21^. FSTL-1 has been implicated in the pathogenesis of pulmonary fibrosis development^22^, emphysema^23^, autoimmunity^24-26^, cardiovascular diseases^27^, and complication associated with Lyme disease^28^. In the context of inflammation, FSTL-1 regulates innate immune cell signaling in macrophages via inflammasome activation and NR4A1 signaling^25,29-31^. Furthermore, FSTL-1 functions as a ligand for the receptor CD14^32^. Despite these observations, how FSTL-1 contributes to antibacterial host defense and the cellular and molecular constituents of FSTL-1 mediated antibacterial function remains to be elucidated.

Previously, we reported that FSTL-1 influences pulmonary immunity against *K. pneumoniae*^23^ using the FSTL-1 hypomorphic (Hypo) mouse model, which exhibited reduced FSTL-1 expression and significantly elevated IL-17A-producing γδT cells with and without infection^23^. In addition, we also observed that FSTL-1 Hypo mice develop spontaneous emphysema, making a clear assessment of the contribution of FSTL-1 to pulmonary host defense difficult^23^. To circumvent these developmental confounding factors of the FSTL-1 Hypo model, we have generated an inducible, global *Fstl1* knockout mouse model, enabling temporal control of post-natal *Fstl1* deletion.

Using this improved model, we sought to test whether FSTL-1 promotes pulmonary immunity against *K. pneumoniae*, as well as the cellular and molecular constituents of FSTL-1 pulmonary immune signaling. The current studies identified that 1) FSTL-1 is induced in the lung following bacterial infection, 2) that FSTL-1 is necessary and sufficient for bacterial immunity and 3) FSTL-1-dependent immunity requires the presence of neutrophils as well as CD14 and NR4A1 expression. Together, these findings establish that FSTL-1 drives neutrophil-mediated bacterial immunity in a CD14- and NR4A1-dependent manner, suggesting that FSTL-1 is a lung tissue-derived signal that modulates lung neutrophil function, representing a possible novel therapeutic target for treating bacterial pneumonia treatment.

## Methods

### Human Subjects/Samples

Bronchoalveolar lavage fluid (BAL) was collected from patients intubated for clinical indications as part of an approved observational study (IRB #STUDY20040201). De-identified clinical data was acquired from the hospital electronic health record to assess microbiological data. Specifically, respiratory culture results including the presence of white blood cells (WBCs), and bacteria on gram stain, and the growth of bacterial isolates in standard culture. Patients with both the presence of “moderate” or “many” WBCs and bacteria on gram stain as well as growth of a potential pathogen were classified as “acute infection”; patients with “no” or “rare” WBCs and no bacterial growth on culture were classified as “healthy controls”.

### Animals

All animal protocols were approved by the University of Pittsburgh Institutional Animal Care and Use Committee (IACUC, protocols #25066905 and #23104015), and all experiments were conducted in accordance with the guidelines and regulations set forth in the Animal Welfare Act (AWA) and PHS Policy on Humane Care and Use of Laboratory Animals. Age- and sex-matched male and female mice between 8-12 weeks old at the beginning of experiments were used for this study.

Wild-type (C57BL/6J, JAX: 00064), *Cd14*^*-/-*^ (B6.129S4-*Cd14*^*tm1Frm*^/J, JAX: 003726)^33^, *Nr4a1*^*-/-*^ (B6;129S2-*Nr4a1*^*tm1Jmi*^/J, JAX: 006187)^34^, Cre-ERT2 (B6.129-*Gt(ROSA)26Sor*^*tm1(cre/ERT2)Tyj*^/J, JAX:008463)^35^, MRP8-Cre (B6.Cg-Tg(S100A8-cre,-EGFP)1llw/J, JAX: 021614)^36^ mice were purchased from the Jackson Laboratory. The *Fstl1-*floxed (*Fstl1*^*fl/fl*^) mice^37^ were a generous gift from the laboratory of Dr. Raphael Hirsch and the ROSA-iDTR mice^38^ were a generous gift from the laboratory of Dr. George Gittes.

Tamoxifen-inducible *Fstl1*^*-/-*^ strain was generated by crossing the Cre-ERT2^35^ with *Fstl1*^*fl/fl*^ knockout mice^37^ to yield the global, tamoxifen-inducible *Fstl1*^*-/-*^mice. Cre-negative littermate controls and *Fstl1*^*-/-*^ mice were treated with 4 mg tamoxifen via intraperitoneal injection (IP) total of three times (two consecutive days, one day off, and the last dose). Experiments were performed at least 40 days post initial tamoxifen administration.

Diphtheria-toxin (DT) inducible *Mrp8*-Cre transgenic mice (PMN^DTR^), in whom diphtheria toxin (DT) treatment readily depletes mature neutrophils, were generated by crossing MRP8-Cre^36^ and ROSA-iDTR^38^ as described previously^39^. *Mrp8*-Cre positive mice were treated with 250 ng vehicle or DT via IP infection 24 hours prior to and on the day of infection.

### *Klebsiella pneumoniae* infection

*In vivo* infection of *K. pneumoniae 43816* (*Kp43816*), a hypervirulent mucoid strain^40^, was performed via oropharyngeal injection (o.p.) as described previously^41,42^. Each animal received *Kp43816* at 5,000–10,000 CFU, except for *Cd14*^*-/-*^ animals who received reduced dosed of 500–1,500 CFU to ensure survival of the animals for the duration of the experiment.

### Bacterial burden analysis

Left lung lobes and spleens of experimental mice were separately harvested into 1 ml PBS and homogenized. Homogenized cell suspensions were diluted in tryptic soy broth (TSB) and bacterial burden was quantified by liquid broth turbidimetry as previously described^43^.

### Quantitative PCR

Right upper and middle lung lobes of experimental mice were processed for qPCR as described previously^41^. See Table S1 for primer assay information.

### Flow cytometry

Right bottom lung lobes of experimental mice were processed and stained for flow cytometry as described previously^41^. See Table S2 for antibody information.

### Enzyme-linked immunosorbent assay (ELISA)

FSTL-1 levels in human BAL fluid and murine lung and serum were quantified using the ELISA kits (R&D) per manufacturer’s instructions.

## Results

### *K. pneumoniae* infection induces pulmonary FSTL-1 expression

We first sought to understand whether FSTL-1 expression was perturbed in the human lung tissue during infection. Patients with acute lung infections had significantly higher FSTL-1 levels in bronchoalveolar lavage (BAL) fluid compared to healthy controls (Figures 1A). We then turned to an *in vivo* model to examine whether lung FSTL-1 levels were similarly altered in wild-type mice during *K. pneumoniae* pulmonary infection. Following lung infection, both FSTL-1 protein and *Fstl1* transcript levels were significantly increased in the lungs (Figures 1B and 1D) of *K. pneumoniae-*infected mice. Notably, serum FSTL-1 protein levels remained unchanged following infection (Figure 1C), suggesting that FSTL-1 induction occurs predominantly within the infected lung.

**Figure 1:**
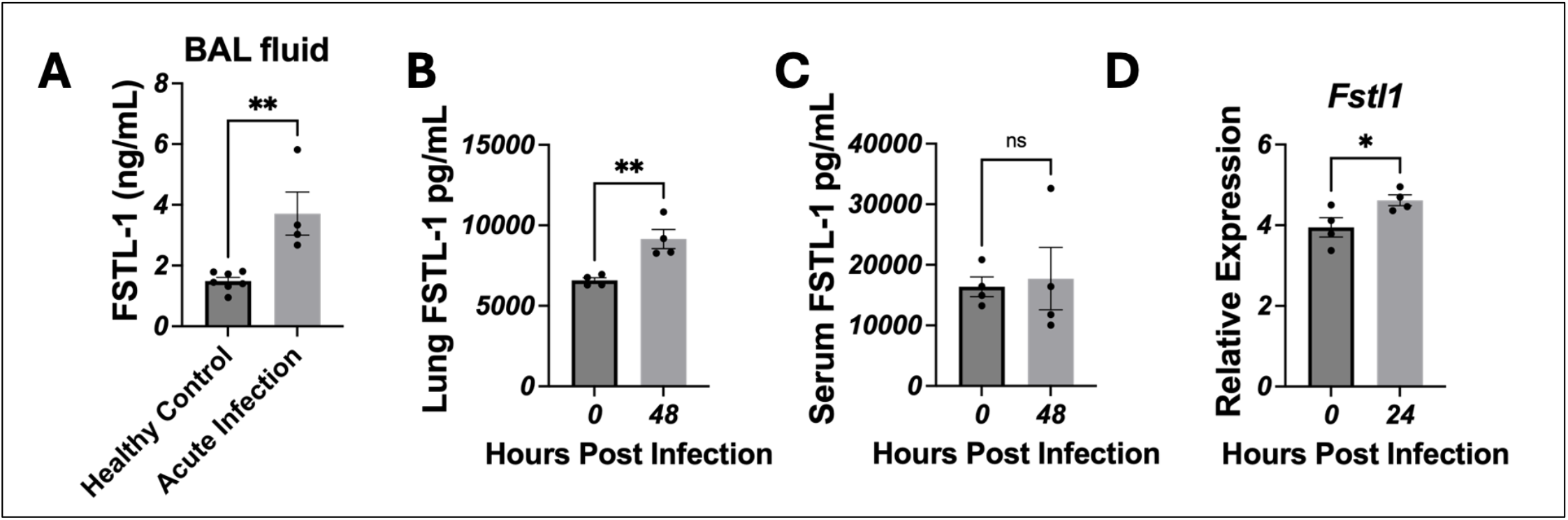
Lung FSTL-1 level is increased by bacterial infection. (A) FSTL-1 protein levels in the BAL fluid of healthy control and patients with acute lung infections were compared by ELISA. WT C57/BL6 mice were infected with *K. pneumoniae* intratracheally (IT). FSTL-1 protein levels in lungs (B) and serum (C) were quantified by ELISA at 0 and 48 hours post *K. pneumoniae* infection. D) WT C57/BL6 mice were infected with *K. pneumoniae* via IT. FSTL-1 transcript level in lungs of WT C57/BL6 mice were quantified by qPCR at 0 and 24 hours post *K. pneumoniae* infection. n= 4-7 (A), 4 (B-D), mean ± SEM., comparison by unpaired *t*-test where ns: not significant, *: p < 0.05, and **: p < 0.01.

### FSTL-1 is required for pulmonary bacterial clearance and neutrophil recruitment during *K. pneumoniae* infection

To determine how FSTL-1 contributes to pulmonary host defense, we employed an inducible *Fstl1*^*-/-*^ mouse model generated by crossing the tamoxifen-inducible Cre recombinase transgenic mouse, B6.129-Gt(ROSA)26Sor^tm1(cre/ERT2)Tyj^/J (Jackson Laboratories #:008463) strain^35^, with germline *Fstl1*^*fl/fl*^ mice^37^. This strategy improved upon our prior work evaluating the role of FSTL-1 during *K. pneumoniae* lung infection using the FSTL-1 hypomorphic (Hypo) mouse model^41^, and represents a significant improvement in three key ways:

1) the inducible *Fstl1*^*-/-*^ model avoids the critical phenotype of spontaneous pulmonary emphysema^23^, 2) *Fstl1* deletion in adulthood circumvents the profound lung development defect associated with germline FSTL-1 deletion, which is lethal, and importantly, 3) the inducible *Fstl1*^*-/-*^ mice does not have aberrant homeostatic Type-17 immune function as evidenced by comparable frequencies of IL-17A-producing γδT cells to WT mice (Figure S1), an immune phenotype that likely influenced our prior observations in the FSTL-1 Hypo mouse^41^.

Using the inducible FSTL-1 knockout, *Fstl1*^*-/-*^ mice exhibited significantly increased lung bacterial burden compared to litter mate controls following *K. pneumoniae* pulmonary infection (Figure 2A). *Fstl1*^*-/-*^ mice also had impaired pulmonary neutrophil recruitment and reduced expression of inflammatory cytokines and chemokines that mediate neutrophil recruitment (Figures 2B-G). These findings indicate that endogenous FSTL-1 promotes pulmonary bacterial control and the recruitment of neutrophils following *K. pneumoniae* infection.

**Figure 2:**
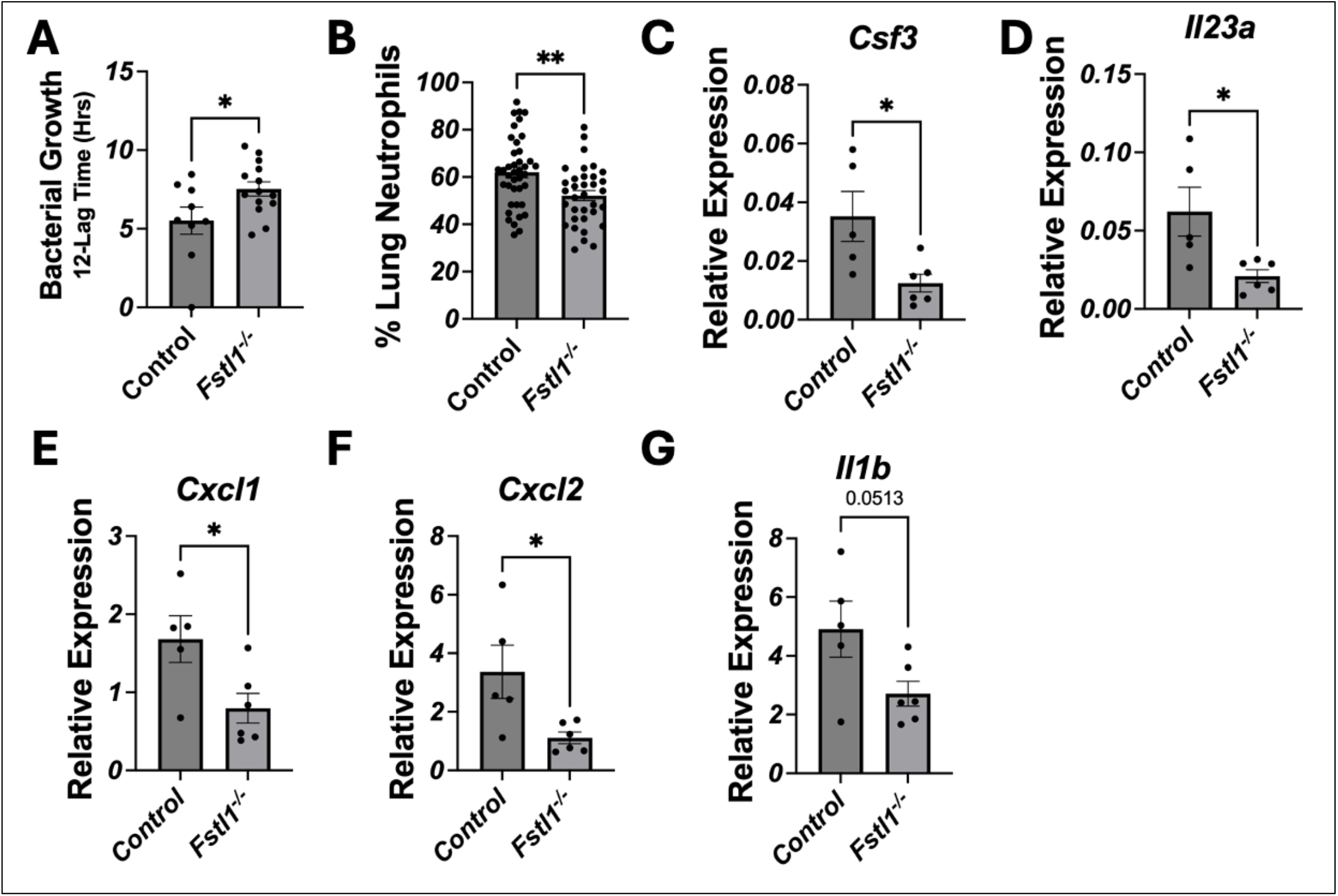
FSTL-1 is essential in controlling *K. pneumoniae* lung burden and mediates neutrophil recruitment and neutrophil-associated cytokines/chemokines. *Fstl1*^*-/-*^ mice and litter mate controls were infected with *K. pneumoniae* o.p. A) Lung bacterial burden was quantified 72 hours post infection via turbidimetry. B) Lung neutrophil (CD11b^+^Gr-1^+^) abundance was quantified 48 hours post infection via flow cytometry. C-G) Lung gene expression was quantified 24 hours post infection by qRT-PCR. n= 9-14 (A), 34-40 (B), 5-6 (C-G), mean ± SEM., comparison by unpaired *t*-test where *: p < 0.05, and **: p < 0.01. Select p values are shown.

### Recombinant FSTL-1 enhances pulmonary host defense in a neutrophil-dependent manner

To determine whether the susceptibility of *Fstl1*^*-/-*^ mice to pulmonary bacterial infection was specifically due to depletion of FSTL-1, we sought to rescue the defect by administering recombinant FSTL-1 (rFSTL-1). We found that rFSTL-1 promoted bacterial clearance in *Fstl1*^*-/-*^ mice (Figure 3A). We then tested whether rFSTL-1 could similarly enhance host defense in WT animals (Figure 3). rFSTL-1 treatment significantly reduced lung bacterial burden and increased pulmonary neutrophil abundance following *K. pneumoniae* infection (Figure 3B-C), suggesting that FSTL-1-mediated antibacterial immunity was associated with neutrophils. To determine whether neutrophils were required for the protective effects of rFSTL-1, we used two independent models of neutrophil depletion. In both antibody-mediated neutrophil depletion and the diphtheria toxin-inducible genetic neutrophil depletion (*Mrp8-Cre*;iDTR mice) approach, rFSTL-1 failed to improve pulmonary bacterial control in the absence of neutrophils (Figure 3D-E). These findings demonstrate that the protective effect of rFSTL-1 against *K. pneumoniae* requires neutrophils.

**Figure 3:**
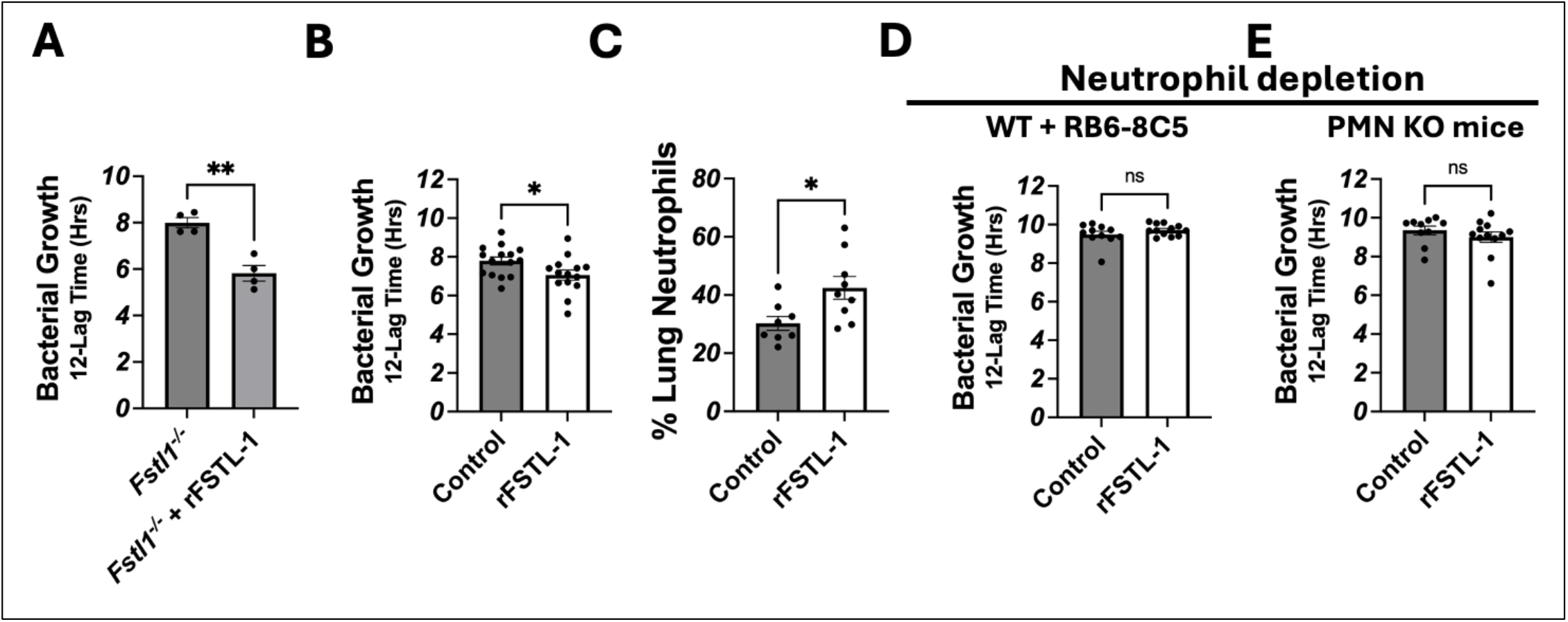
Recombinant FSTL-1 protects against *K. pneumoniae* infection in a neutrophil dependent manner. *Fstl1*^*-/-*^ mice (A) or WT C57/BL6 mice (B-C) were infected with *K. pneumoniae* o.p. Animals received 800 ng vehicle or recombinant FSTL-1 via IP injection at 0- and 24-hours post infection. A-B) Lung bacterial burden was quantified 72 (A) or 48 (B) hours post infection by turbidimetry. C) Lung neutrophil recruitment was quantified by flow cytometry at 12 hours post infection. D-E) Effect of rFSTL-1 treatment on lung bacterial burdens in two neutrophil depletion models were investigated. *K. pneumoniae* infection, rFSTL-1 treatment, and lung bacterial burden quantifications were performed as described in panel A. D) WT C57/BL6 animals were treated with vehicle or RB6-8C5 via IP injection 24 hours prior to infection. E). Diphtheria toxin (DT)-inducible *PMN*^*DTR*^ mice were treated with 250 ng vehicle or DT via IP injection 24 hours prior to and on the day of infection. n = 4 (A), 14-15 (B), 8-9 (C), 11-12 (D) or 10-12 (E). mean ± SEM., comparison by unpaired *t*-test where ns: not significant and *: p < 0.05.

### FSTL-1-mediated protection requires CD14 and NR4A1

Circulating neutrophils undergo tissue-specific functional adaptation in the lung, associated with induction of a transcriptional program characterized by expression of *Cd14* and *Nr4a1*, genes identified to mediate pulmonary immunity against infections by Gram negative pathogens, which had been previously shown to mediate FSTL-1-dependent signaling^*16,17,42,44*^. We thus tested whether CD14 or NR4A1 were required for FSTL-1 mediated antibacterial defense. Administration of rFSTL-1 failed to reduce lung bacterial burden or improve survival in *Cd14*^*-/-*^ mice (Figure 4A-B). Similarly, rFSTL-1 also failed to improve pulmonary bacterial control or survival in *Nr4a1*^*-/-*^ mice (Figure 4C-D). Together, these findings indicate that the protective effect of FSTL-1 in *K. pneumoniae* lung infection requires CD14- and NR4A1-dependent pathways.

**Figure 4:**
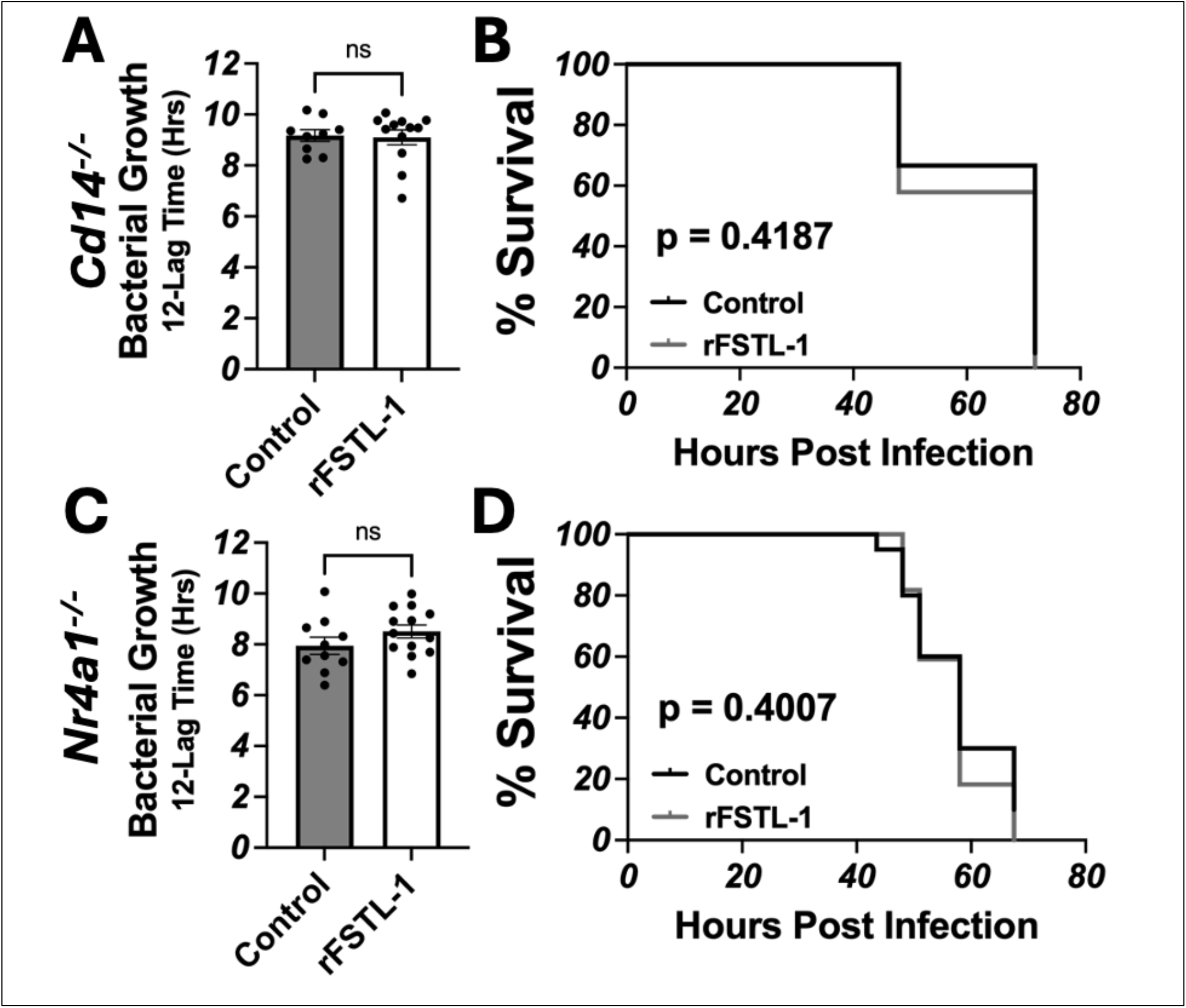
FSTL-1 dependent protection requires *Cd14* and *Nr4a1*. Animals were infected with 1,000 CFU (*Cd14*^*-/-*^) or 5,000 CFU (*Nr4a1*^*-/-*^) of *K. pneumoniae* via o.p. and received 800 ng recombinant FSTL-1 or vehicle at 0 and 24 hours post infection. Lung bacterial burden of *Cd14*^*-/-*^ (A) or *Nr4a1*^*-/-*^ (C) mice were assessed at 48 hours post infection by turbidimetry. Survival of *Cd14*^*-/-*^ (B) or *Nr4a1*^*-/-*^ (D) mice who received vehicle or 800 ng recombinant FSTL-1 at 0 and 24 hours post infection. n = 19-22 (A, C) or 9-13 (B, D). Comparison by t-test (A, C) or Mantel-Cox test (B, D). Ns: no statistical difference. Select p values are shown.

## Discussion

### FSTL-1 is a marker of lung inflammation and mediates pulmonary host defense

Follistatin-like 1 (FSTL-1) is a secreted glycoprotein that has been implicated in a broad range of inflammatory and tissue-remodeling processes^29-31^. Increased FSTL-1 levels in tissues, serum, and synovial fluid have been reported in inflammatory diseases including rheumatoid arthritis and osteoarthritis and have been associated with increased disease activity and tissue inflammation^20,21,25^. FSTL-1 has also been investigated in other inflammatory and vascular diseases in which its biological effects appear to be highly dependent on cellular context and tissue environment^29,30,45^. These observations have established FSTL-1 as a biomarker as well as mediator of inflammatory diseases. Importantly, the present study identifies FSTL-1 as a biomarker of inflammation in bacterial lung disease (bacterial infection) and identifies the cellular and molecular mechanisms by which FSTL-1 mediates protection against pulmonary bacterial infection.

FSTL-1 is expressed in multiple mesenchymal cell populations and has important functions in lung development and tissue remodeling^19,23,46,47^. During lung development, FSTL-1 regulates BMP4 signaling and contributes to normal pulmonary morphogenesis^19^. In the adult lung, FSTL-1 has been implicated in fibroblast activation and pulmonary fibrosis, in part through modulation of TGFβ signaling^22^. Studies of experimental lung injury have demonstrated that increased FSTL-1 can promote inflammatory and fibrotic responses, whereas genetic or antibody-mediated depletion of FSTL-1 can attenuate fibrosis in various models^22,24,48^. Thus, previous studies have primarily considered FSTL-1 in the context of lung development, tissue injury and pathological remodeling.

Irrespective of the cellular origin, FSTL-1 can influence inflammatory responses through interactions with immune cells. FSTL-1 has been reported to mediate macrophage and monocyte responses, including NLRP3 inflammasome activation and IL-1β production^31^. FSTL-1 also acts on alveolar macrophages to suppress LPS-induced inflammation in an NR4A1-dependent manner, providing evidence that FSTL-1 regulates immune-cell responses^30,49,50^.

Our previous studies have provided evidence that FSTL-1 contributes to pulmonary immunity during *K. pneumoniae* infection by using an FSTL-1 hypomorphic (Hypo) model^23^. The present study improves upon these observations using a superior inducible *Fstl1*^*-/-*^ model to more directly examine the requirement for FSTL-1 during infection in adult animals. Inducible *Fstl1* deletion circumvents the confounding effects of 1) germline FSTL-1 deficiency on development or immune-system maturation, 2) spontaneous pulmonary emphysema development, and 3) enhanced homeostatic type-17 immune response. This distinction is relevant because γδT cell-derived IL-17A substantially influence pulmonary antibacterial immunity and bacterial burden^23^. Thus, the current model provides an opportunity to more precisely examine the contribution of FSTL-1 to antibacterial host defense.

This study shows that FSTL-1 is induced locally in the lung during *K. pneumoniae* pneumonia and contributes functionally to pulmonary host defense. FSTL-1 is elevated in BAL fluid from patients with acute lung infections, and both FSTL-1 protein and *Fstl1* transcript are increased in the lungs of mice following *K. pneumoniae* infection. Notably, FSTL-1 protein level was not significantly increased in the serum of infected mice, suggesting that its induction during pneumonia is driven by lung tissue expression as a localized response to infection rather than as a generalized systemic response. Confirming a functional role, inducible *Fstl1* deletion resulted in increased pulmonary bacterial burden and impaired neutrophil recruitment to the lungs. Conversely, administration of recombinant FSTL-1 (rFSTL-1) restored bacterial control in FSTL-1-deficient mice and further enhanced bacterial clearance in WT mice as well. Together, these findings extend previous observations linking FSTL-1 to pulmonary inflammation and demonstrate that FSTL-1 is not merely associated with inflammation in lung disease but actively contributes to antibacterial host defense. Furthermore, our data establish that neutrophils are an important cellular mediator through which FSTL-1 promotes host defense. These findings suggest that FSTL-1 may function within the lung microenvironment to promote the recruitment and/ or functional adaptation of neutrophils during bacterial infection.

### FSTL-1 as a potential host-directed therapeutic

The ability of exogenous FSTL-1 to enhance bacterial clearance raises the possibility that FSTL-1 could be utilized as part of a host-directed therapeutic strategy. Previous studies have demonstrated beneficial effects of recombinant FSTL-1 in tissue repair and regeneration, including cardiac regeneration following myocardial injury^51^ and would healing^52^. These support the rationale and feasibility of therapeutically modulating extracellular FSTL-1 activity. In the present study, administration of rFSTL-1 during *K. pneumoniae* pneumonia reduced pulmonary bacterial burden in WT mice and restored bacterial control in FSTL-1 deficient animals.

The observed therapeutic effect of FSTL-1 is particularly notable for the requirement of neutrophils, as rFSTL-1 failed to improve bacterial clearance in neutrophil depleted mice. Thus, FSTL-1-mediated antibacterial protection acts to augment host immune function rather than as an antimicrobial. This finding may be particularly relevant for antibiotic-resistant infections which may otherwise be limited in efficacy by antimicrobial resistance; restoration or enhancement of host innate immunity could complement antimicrobial therapy.

Several questions remain before FSTL-1 can be considered as a viable therapeutic strategy. First, the receptor and proximal signaling pathways through which extracellular FSTL-1 acts on lung neutrophils remain incompletely evaluated. Though our findings implicate an FSTL-1/CD14/NR4A1 signaling axis in this context, several FSTL-1-interacting proteins and signaling pathways have been identified in other cellular contexts^30,53^, warranting further investigation of FSTL-1 relevant receptor(s) and downstream mechanisms in lung neutrophils during bacterial pneumonia^41,54^. Second, the optimal dose, route, timing, and duration of FSTL-1 administration remain undetermined. Finally, because FSTL-1 has context-dependent effects on inflammation, tissue remodeling, and fibrosis, the consequences and potential adverse effects of sustained or repeated FSTL-1 administration will require careful evaluation. Our findings therefore provide proof of principle for FSTL-1 as a host-directed modulator of antibacterial immunity while establishing a framework for future studies to define its therapeutic paremeters and mechanism of action.

### FSTL-1 is a constituent of a lung-adapted neutrophil program

Neutrophils have been historically viewed as relatively homogeneous circulating effector cells that respond to tissue infection primarily through canonical antimicrobial functions^55,56^. Contemporary evidence instead demonstrates that neutrophils acquire tissue-specific transcriptional and functional states in response to local environmental cues^16,17^. Ballesteros *et al*. have demonstrated that tissue environments can instruct distinct neutrophil transcriptional states, including a lung-associated program characterized by expression of genes such as *Cd14* and *Nr4a1*^16^. Subsequent work has reinforced the concept that neutrophils exist on a dynamic continuum shaped by tissue-specific signals rather than as a single uniform population^17^.

Our findings suggest that FSTL-1 may be one component of the local lung environment that participates in this process. The protective effect of rFSTL-1 was absent in both *Cd14*- and *Nr4a1*-deficient mice. These findings place CD14 and NR4A1 downstream of, or functionally required for, FSTL-1-mediated antibacterial immunity. Together, these observations suggest that FSTL-1 may contribute not simply to the recruitment of neutrophils into the infected lung, but also to their functional adaptation after arrival.

The role of CD14 in neutrophil biology is less clearly defined than its established functions in monocytes and macrophages, and studies of CD14 in pulmonary infection have produced context-dependent findings^57-61^. Nevertheless, the identification of *Cd14* as part of the lung-associated neutrophil transcriptional program and the loss of FSTL-1-mediated protection in *Cd14*-deficient mice suggest that CD14 represents an important component of the FSTL-1-responsive neutrophil state. Similarly, the requirement for NR4A1 is consistent with the broader role of this transcription factor in tissue-adapted neutrophil function and pulmonary host defense^42^.

An important unresolved question is how local tissue cells communicate with circulating neutrophils to establish these tissue-specific states. The lung contains abundant populations of fibroblasts, endothelial cells, other stromal cells and tissue-resident immune cells capable of producing extracellular mediators that can influence immune-cell behavior^62-67^. FSTL-1 is expressed predominantly by fibroblast and endothelial cell populations rather than by hematopoietic cells, providing a potential mechanism for communication from the lung microenvironment to recruited neutrophils^23,53,68^. Indeed, FSTL-1 has been described as a product of mesenchymal cells and can act on immune cells in inflammatory tissues^30,31,45,69^.

Thus, our findings support a conceptual model in which infection-induced FSTL-1 production by lung stromal cells contributes to the recruitment and functional lung-specific adaptation of neutrophils, which then supports bacterial clearance in a CD14- and NR4A1-dependent manner. This model raises several important questions for future investigation, including which lung-resident cell population is the dominant source of FSTL-1 during infection, how neutrophils receive FSTL-1-dependent signals, and whether FSTL-1 directly regulates the acquisition of the lung-specific neutrophil transcriptional program or acts indirectly through other inflammatory mediators. Defining these mechanisms will help clarify how tissue-derived signals shape neutrophil function during infection and may reveal additional opportunities for therapeutic modulation of antibacterial host defense.

In summary, our study identifies FSTL-1 as an inducible component of the pulmonary inflammatory response that is required for effective host defense against *K. pneumoniae*. FSTL-1 promotes neutrophil recruitment and neutrophil-mediated antibacterial protection CD14- and NR4A1-dependent pathways. These findings expand the biological functions of FSTL-1 in the lung beyond inflammation and tissue remodeling and identify a previously unrecognized connection between a lung-derived extracellular mediator and tissue-adapted neutrophil immunity.

## Acknowledgements

This research was supported by NIH R01HL158576, K08HL128809 and UPMC Children’s Hospital of Pittsburgh (BTC).

We thank Dr. Taylor Eddens for insightful discussion. We are grateful to the Hirsch and the Gittes laboratories for providing key mice strains. We also thank the Flow Cytometry Core at the UPMC Children’s Hospital of Pittsburgh (RRID:SCR_025223) for cytometer access, and the Division of Laboratory Animal Resources (DLAR) for animal care.

## Author Contribution

NS, MH, and BTC designed experiments. NS, MH, BTC, JP, AA and TW executed experiments and analyzed data. NS drafted the manuscript. MH and BTC edited the manuscript. NS, MH, JP, AA, TW, NWF, AD and BTC reviewed and approved the final manuscript.

