## Supplementary Data 1 for "Follistatin-like 1 Promotes Neutrophil-mediated Host Defense against *Klebsiella pneumoniae*"

### Supplementary Tables

| Gene | Source | Identifier |
|---|---|---|
| Fstl1 (mouse) | ThermoFisher Scientific | Mm00433371_m1 |
| Csf3 (mouse) | ThermoFisher Scientific | Mm00438334_m1 |
| Il23a (mouse) | ThermoFisher Scientific | Mm00518984_m1 |
| Cxcl1 (mouse) | ThermoFisher Scientific | Mm04207460_m1 |
| Cxcl2 (mouse) | ThermoFisher Scientific | Mm00436450_m1 |
| Il1b (mouse) | ThermoFisher Scientific | Mm00434228_m1 |
| Hprt (mouse) | ThermoFisher Scientific | Mm03024075_m1 |

**Table S1: qPCR primer assays**

| Antibody | Clone | Source | Identifier |
|---|---|---|---|
| APC anti-mouse/ human CD11b | M1/70 | BioLegend | 101212 |
| FITC anti-mouse Ly6G | 1A8 | BioLegend | 127606 |
| FITC anti-mouse Gr-1 | RB6-8C5 | BioLegend | 108406 |
| BV786 anti-mouse CD11c | HL3 | BD | 563735 |
| PerCP-Cy5.5 anti-mouse CD45 | 30-F11 | BD | 561869 |
| FITC anti-mouse IL17-A | eBio17B7 | Invitrogen | 11-7177-81 |
| PerCP-Cy5.5 anti-mouse IL17-A | eBio17B7 | eBioscience | 45-7177-82 |
| PE anti-mouse $\gamma\boldsymbol{\delta}$TCR | eBioGL3 | eBioscience | 12-5711-82 |
| APC-Cy7 anti-mouse CD3e | 17A2 | BD | 560590 |
| ant-mouse CD16/ CD32 | 93 | Invitrogen | 14-0161-85 |
| **Live/dead dye** | **Clone** | **Source** | **Identifier** |
| Aqua Live Dead | N/A | ThermoFisher | L34966 |

**Table S2: Antibodies for FACS**

### Supplementary Figures

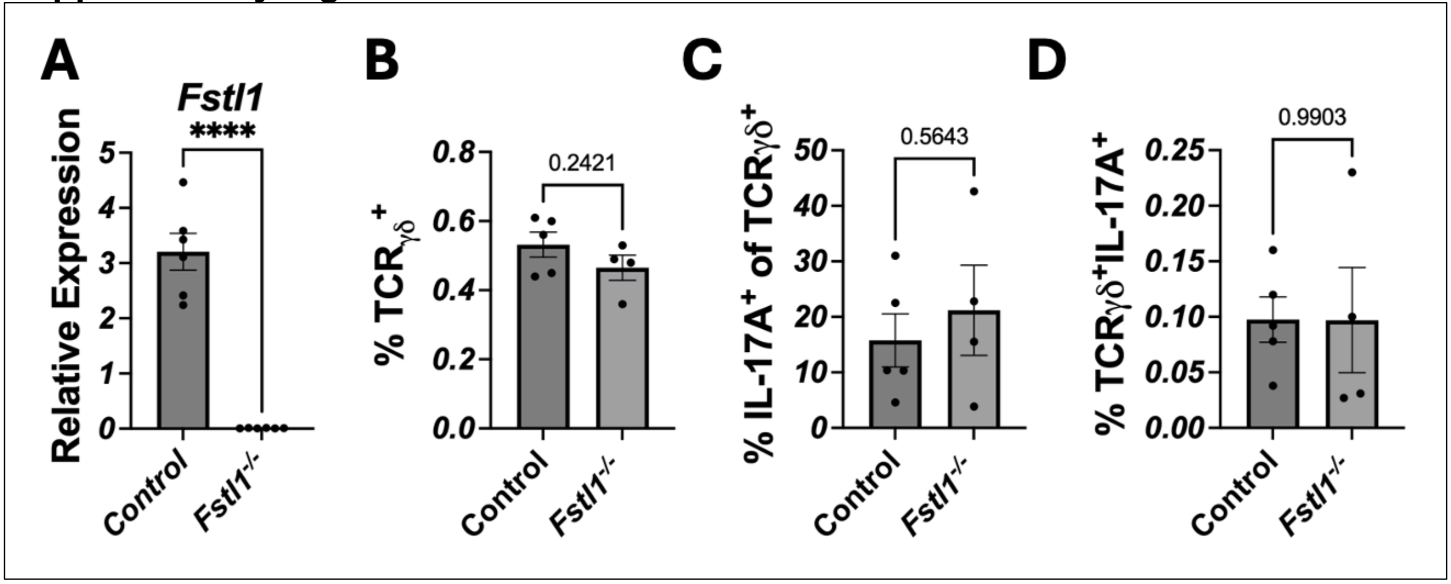


**Figure S1: Inducible *Fstl1*$^{-/-}$ mice have comparable IL-17A-producing γδT cells to the control mice.**

(A) Confirmation of *Fstl1* gene deletion in *Fstl1*$^{-/-}$ mice post tamoxifen treatment. (B-D) Flow cytometry immunophenotyping of lung cells in uninfected, tamoxifen treated litter mate control and *Fstl1*$^{-/-}$ mice. Proportions of γδT cells (B), IL-17A within γδT cells (C), and TCRγδ$^{+}$IL-17A$^{+}$ cells (D) in uninfected animals were analyzed by flow cytometry. n=5-6 (A), 4-5 (B-C), mean ± SEM., comparison by unpaired *t*-test where ****: $p < 0.0001$. Select p values are shown.

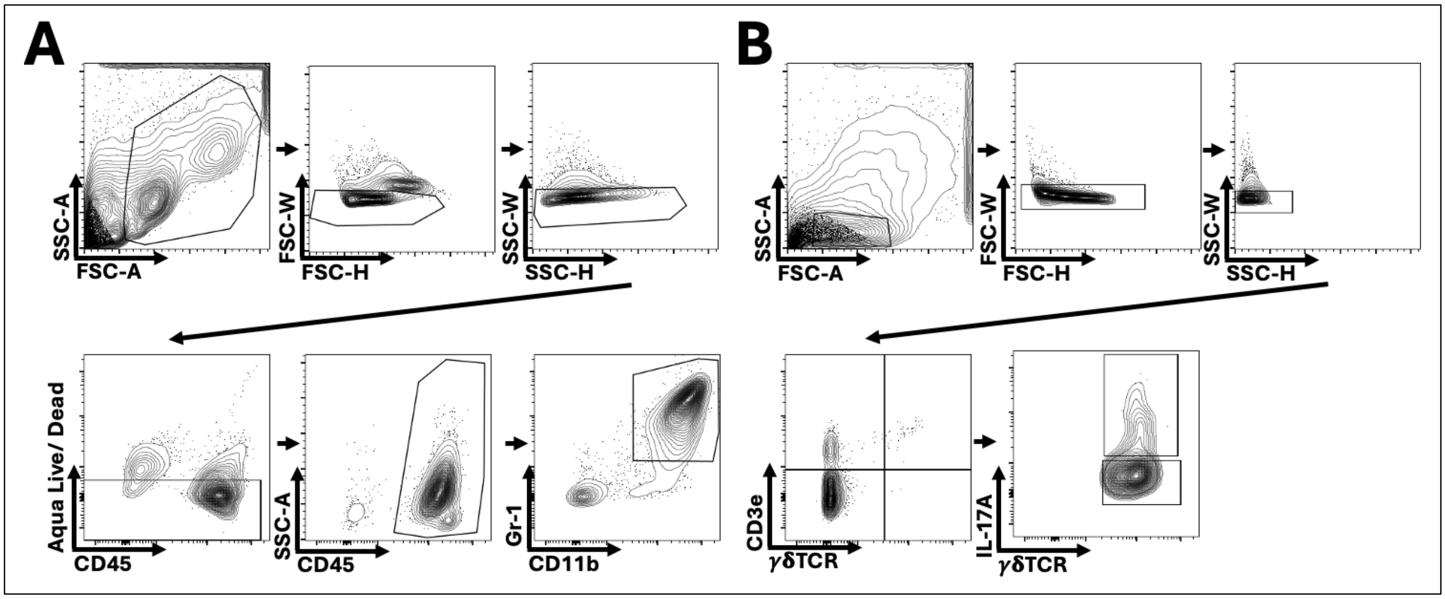


**Figure S2: FACS gating strategy.**
(A) Lung neutrophil phenotyping in Figures 2B and 3C. (B) IL17-A producing γδT cell phenotyping in Figure S1.
